# T9GPred: A Comprehensive Computational Tool for the Prediction of Type 9 Secretion System, Gliding Motility and the Associated Secreted Proteins

**DOI:** 10.1101/2023.03.31.535141

**Authors:** Ajaya Kumar Sahoo, R. P. Vivek-Ananth, Nikhil Chivukula, Shri Vishalini Rajaram, Karthikeyan Mohanraj, Devanshi Khare, Celin Acharya, Areejit Samal

**Author notes:** Corresponding author (A. Samal). **Address for correspondence:** Areejit Samal Computational Biology Group, The Institute of Mathematical Sciences (IMSc), CIT Campus, Taramani, Chennai 600113 India.

## Abstract

Type 9 secretion system (T9SS) is one of the least characterized secretion systems exclusively found in the *Bacteroidetes* phylum which comprise various environmental and economically relevant bacteria. While T9SS plays a central role in bacterial movement termed gliding motility, survival and pathogenicity, there is an unmet need for a comprehensive tool that predicts T9SS, gliding motility and proteins secreted via T9SS. In this study, we develop such a computational tool, Type 9 secretion system and Gliding motility Prediction (T9GPred). To build this tool, we manually curated published experimental evidence and identified mandatory components for T9SS and gliding motility prediction. We also compiled experimentally characterized proteins secreted via T9SS and determined the presence of three unique types of C-terminal domain signals, and these insights were leveraged to predict proteins secreted via T9SS. Notably, using recently published experimental evidence, we show that T9GPred has high predictive power. Thus, we used T9GPred to predict the presence of T9SS, gliding motility and associated secreted proteins across 693 completely sequenced *Bacteroidetes* strains. T9GPred predicted 402 strains to have T9SS, of which 327 strains are also predicted to exhibit gliding motility. Further, T9GPred also predicted putative secreted proteins for the 402 strains. In a nutshell, T9GPred is a novel computational tool for systems-level prediction of T9SS and streamlining future experimentation. The source code of the computational tool is available in our GitHub repository: https://github.com/asamallab/T9GPred. The tool and its predicted results are compiled in a web server available at: https://cb.imsc.res.in/t9gpred/.

## INTRODUCTION

Type 9 secretion system (T9SS) is one of the recently characterized secretion systems found particularly in members of the *Bacteroidetes* phylum [1–4]. *Bacteroidetes* is a diverse phylum of bacteria found in different environmental niches including animal gut microbiota, where they are either beneficial or pathogenic based on their secretory factors [5,6]. In some of these *Bacteroidetes* strains, T9SS has been experimentally identified to play a central role in movement of the bacteria [1,7–9], assimilation of ions [10], and in the secretion of various polysaccharide degrading enzymes [10,11], adhesins [12] and virulent proteins [2,13]. Additionally, T9SS is found to be essential for the growth and survival of some species [13–15]. Thus, identification of T9SS is imperative to understand the underlying microbial activity.

T9SS was initially characterized in two different *Bacteroidetes* species, *Porphyromonas gingivalis* (human periodontal pathogen) [1,16] and *Flavobacterium johnsoniae* [1]. In particular, McBride and colleagues had extensively characterized the protein components associated with gliding motility of *F. johnsoniae* [17,18] and later found a large overlap with the T9SS components [3]. Additional experiments proved that the gliding motility is facilitated by secretion of adhesins through T9SS [19,20]. Further, several groups have identified functionalities of the core protein components of T9SS namely, GldK, GldL, GldM, GldN and SprA [21–24]. James *et al*. [25] have shown that the GldLM complex is conserved across members of the *Bacteroidetes* phylum and is mediated through proton flow to drive the secretion via T9SS [22,26]. There are other such independent studies focussed on T9SS or gliding motility in members of the *Bacteroidetes* phylum [7,8,27], but there is scarce attempt toward systems-level understanding of the components associated with T9SS or gliding motility.

Secretion via T9SS is a two-step process: (i) transport of protein from cytoplasm to periplasm via Sec export pathway mediated by N-terminal domain; (ii) transport of protein from periplasm to extracellular space via T9SS mediated by C-terminal domain (CTD) [13,28]. Kulkarni *et al.* [29] have experimentally proved that the N-terminal domain and the CTD signal are necessary and sufficient for the secretion of a protein via T9SS. Additionally, they have identified different types of CTD. While efforts have been made to characterize the secreted proteins, there have been limited attempts to identify the CTD types and predict such secreted proteins.

Earlier, Abby *et al.* [4,30] had developed a computational tool, TXSScan to identify different secretion systems including T9SS across bacteria. However, TXSScan is neither specific to T9SS nor does it predict gliding motility or proteins secreted via T9SS. Therefore, in this study, we built a prediction tool which is dedicated to T9SS, gliding motility and the proteins secreted via T9SS. Initially we compiled the proteins associated with T9SS or gliding motility from published literature and generated Hidden Markov Model (HMM) profiles for each protein. By leveraging the compiled experimental knowledge, we identified the key protein components and built a computational tool for prediction of T9SS or gliding motility across members of the *Bacteroidetes* phylum. Note that, we have included gliding motility prediction along with that of T9SS as some *Bacteroidetes* strains having T9SS do not exhibit gliding motility [1,13]. Previously, Veith *et al.* [31] have used HMM based approaches to predict proteins secreted via T9SS. However, their prediction model did not account for different types of CTD. Here we leveraged the different CTD types and their motifs from published literature to generate HMM profiles and built a computational pipeline for the prediction of proteins secreted via T9SS. Finally, we developed a computational tool, **T**ype **9** secretion system and **G**liding motility **Pred**iction (T9GPred), which is accessible at: https://github.com/asamallab/T9GPred. We also compiled the tool and its predictions into a web server namely T9GPred, which is accessible at: https://cb.imsc.res.in/t9gpred/.

## RESULTS

### Identification of 28 protein components associated with T9SS or gliding motility among members of the *Bacteroidetes* phylum

We compiled 23 protein components associated with T9SS (**Table S1**), 22 protein components associated with gliding motility (**Table S2**), and 102 proteins secreted via T9SS (secreted proteins) (**Table S3**) in *Bacteroidetes* strains from published literature (Methods; **Figure 1**). We observed that there is a large overlap of 17 proteins between the proteins associated with T9SS and gliding motility, which is in line with the previous studies reporting conclusive evidence of a link between T9SS and gliding motility in *Bacteroidetes* strains [1,3]. In total, we identified 28 proteins associated with either T9SS or gliding motility among members of the *Bacteroidetes* phylum, which is a much larger set in comparison with the set of 11 proteins used by TXSScan [4,30] for their prediction.

**Figure 1:**
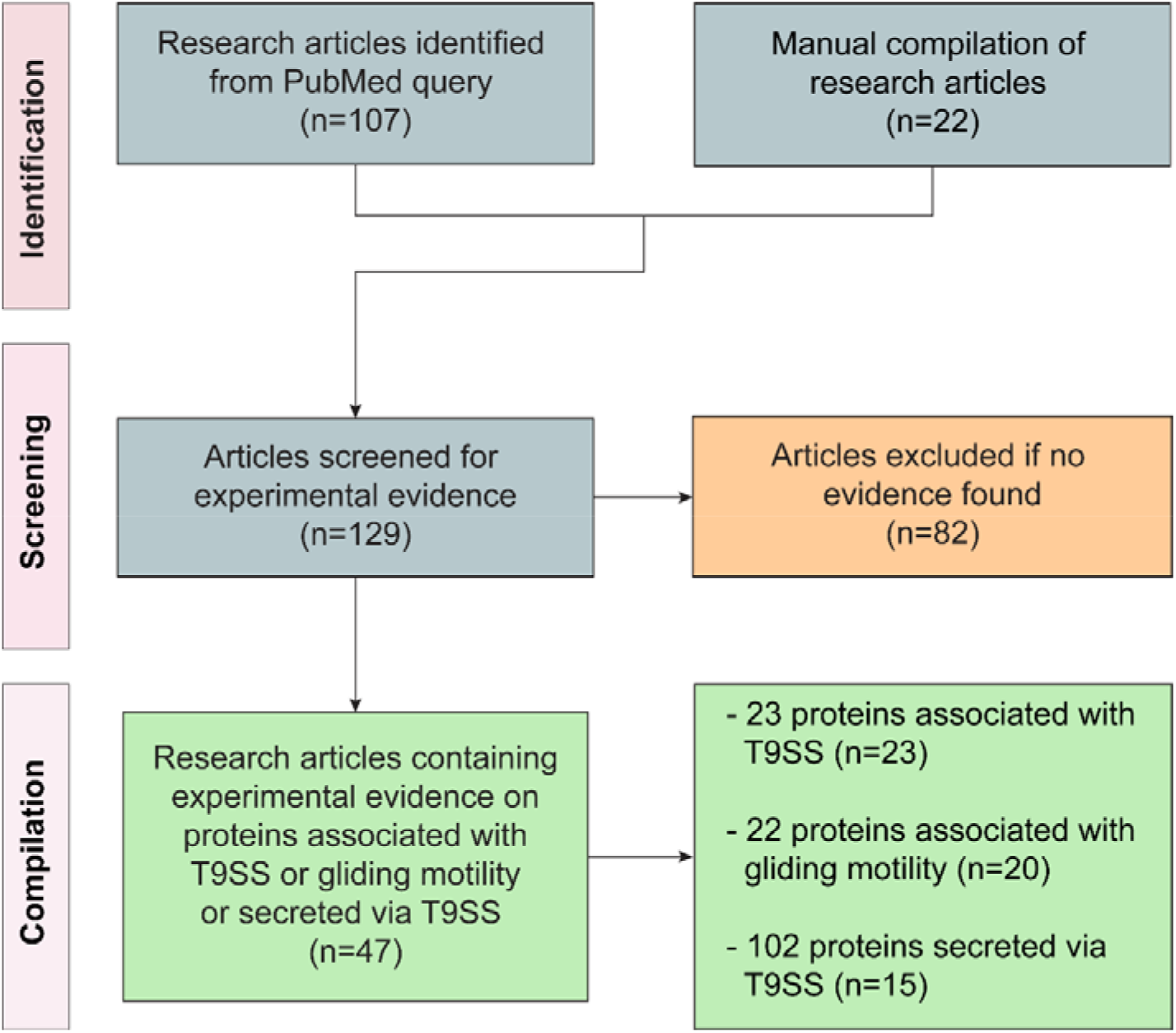
Workflow to identify published research articles containing experimental evidence on the proteins associated with T9SS or gliding motility or proteins secreted via T9SS. This workflow is presented according to the PRISMA statement [44].

### Key protein components of T9SS and gliding motility

During the compilation of associated protein components (**Figure 1**), we observed that not all the components are present in every bacterium of the *Bacteroidetes* phylum that have experimental evidence for T9SS or gliding motility (**Figure 2**). Therefore, we were interested in identifying the key components which are mandatory for a functional T9SS or exhibiting gliding motility. We generated HMM profiles for the 28 identified protein components (Methods; **Supplemental Information**) and checked their presence in 30 *Bacteroidetes* strains that have experimental evidence for the presence of T9SS or gliding motility (**Table S4**). We observed that proteins GldK, GldL, GldM, GldN, SprA, SprE and SprT are always present in the *Bacteroidetes* having T9SS in comparison with those which lack T9SS (**Figure 2**). The proteins GldL and GldM form an inner membrane proton-driven motor that drives protein secretion via T9SS (**Figure 3**) [22,26]. The GldK-GldN ring and SprE facilitate energy transduction from the inner membrane GldLM motor to the outer membrane translocon SprA for the secretion of proteins to the extracellular space (**Figure 3**) [21,22,24,32]. SprT aids in the maturation of the secreted protein but its role in the secretion mechanism is not yet known [16]. Therefore, we designated GldK, GldL, GldM, GldN, SprA and SprE (6 proteins) as mandatory protein components to predict the presence of T9SS.

**Figure 2:**
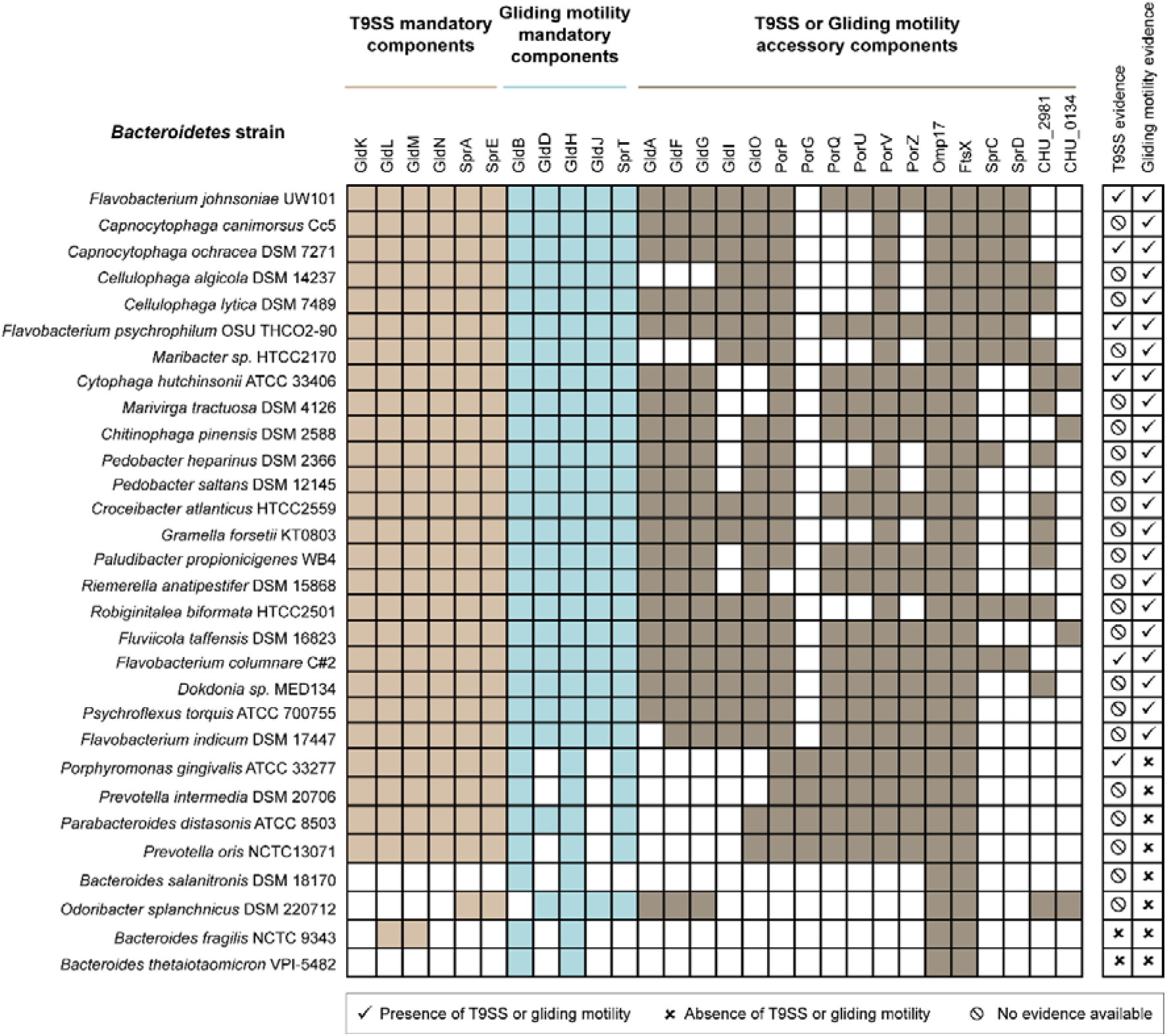
Occurrence of 28 identified protein components associated with T9SS or gliding motility checked across 30 experimentally characterized *Bacteroidetes* strains using the generated HMM profiles. The presence of T9SS mandatory components are denoted by light brown colored box, gliding motility mandatory components are denoted by cyan colored box, and T9SS or gliding motility accessory components are denoted by dark brown colored box. The experimental evidence (presence, absence or no evidence) for T9SS or gliding motility is denoted alongside each *Bacteroidetes* strain.

**Figure 3:**
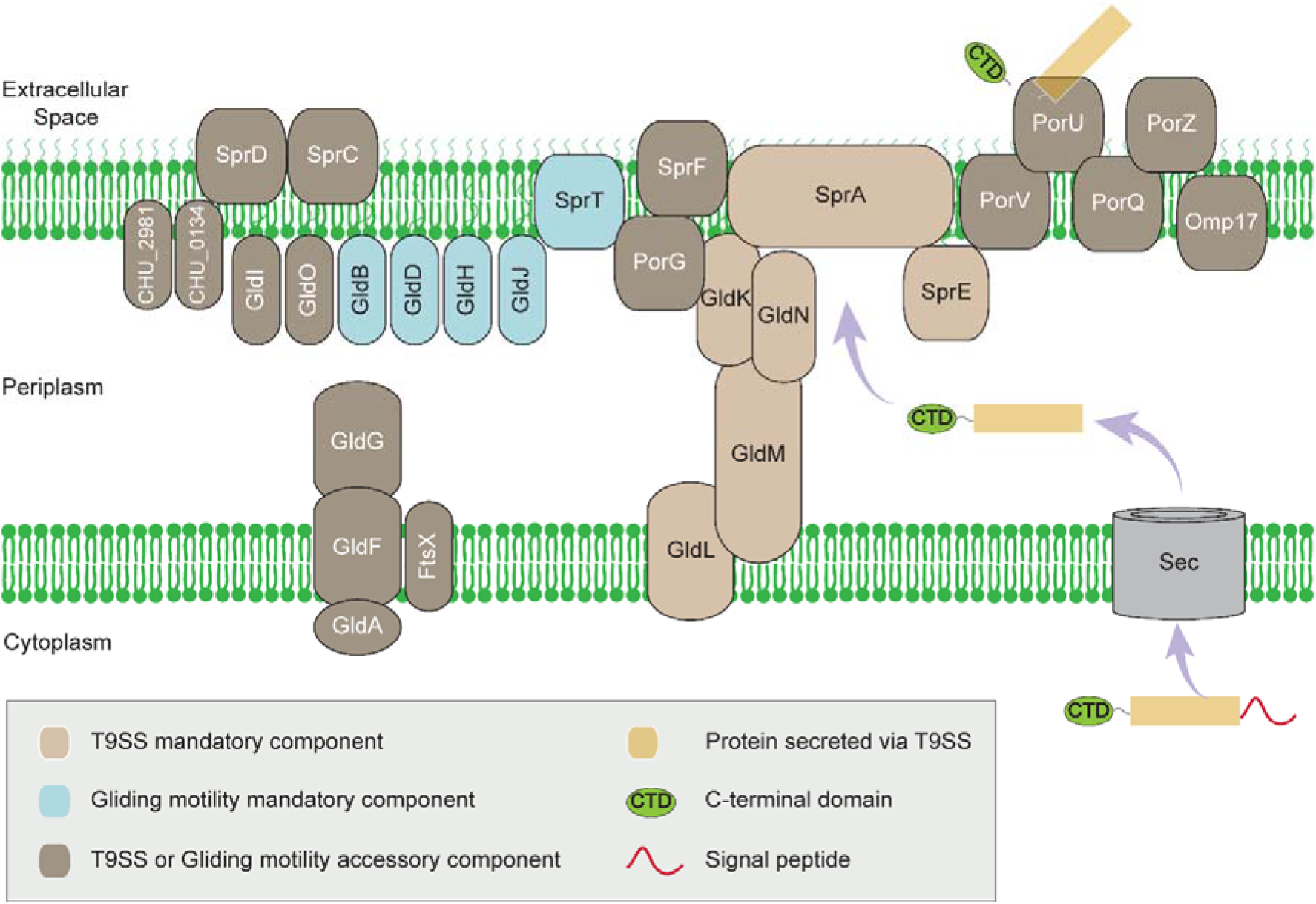
Schematic diagram encapsulating the available information on localization of 28 protein components on the bacterial membrane based on published literature.

In addition to the 6 T9SS mandatory components, we observed that proteins GldB, GldD, GldH, GldJ and SprT are present in the *Bacteroidetes* strains exhibiting gliding motility in comparison with those which lack gliding motility (**Figure 2**). Importantly, McBride *et al.* [3] had earlier characterized the presence of these proteins in *Bacteroidetes* strains exhibiting gliding motility. Therefore, we designated the 6 T9SS mandatory components along with GldB, GldD, GldH, GldJ and SprT (a total of 11 proteins) as mandatory protein components to predict the presence of gliding motility. We designated the remaining 17 proteins as accessory components.

### Computational tool for prediction of T9SS or gliding motility across *Bacteroidetes* **strains**

We leveraged the generated HMM profiles to check for presence of the 28 proteins associated with T9SS or gliding motility in the proteome of 693 completely sequenced *Bacteroidetes* strains (Methods; **Table S5**). If a bacterium contains all 6 T9SS mandatory components in its proteome, we classify it as having T9SS. In addition, if a bacterium contains all the 11 gliding motility mandatory components, we classify it as having gliding motility. Using these criteria, we developed a tool in python programming language which employs the generated HMM profiles for 28 proteins to predict the presence of T9SS or gliding motility across 693 *Bacteroidetes* strains. Our tool identified 402 *Bacteroidetes* strains to have T9SS, of which 327 were found to also have gliding motility. Typically, we observed that 19 of the 28 proteins are present in the 402 *Bacteroidetes* strains predicted to have T9SS (**Figure S1a**), while 22 of the 28 proteins are present in the 327 *Bacteroidetes* strains predicted to have gliding motility (**Figure S1b**). In sum, we developed a computational tool which can predict the presence of both T9SS and gliding motility in members of the *Bacteroidetes* phylum, including mandatory and accessory protein components.

### Validation of T9GPred predictions of T9SS and gliding motility

We validated the T9GPred predictions of T9SS and gliding motility using the recently published experimental evidence (after August 2020) on T9SS and gliding motility that had not been considered to define mandatory components and build T9GPred (**Supplemental Information; Figure S2**). We identified 4 new *Bacteroidetes* strains to have T9SS (**Table S6**) of which all the strains are predicted to have T9SS by T9GPred. Moreover, 2 of the 4 new strains are of new species which were not considered while building the tool for T9SS prediction. We also identified 7 new *Bacteroidetes* strains to have gliding motility (**Table S7**) of which all the strains are predicted to have gliding motility by T9GPred. 4 of 7 new strains having gliding motility are of new species which were not considered while building the tool for gliding motility prediction. Notably, we did not find in the published literature after August 2020, any recently characterized protein components associated with T9SS or gliding motility.

### Classification of the secreted proteins into three C-terminal domain (CTD) types

To classify the proteins secreted via T9SS, we relied on the CTD of the compiled list of 102 secreted proteins with published experimental evidence (Methods). Earlier studies on T9SS have reported the motif sequences for the CTD of the secreted proteins [31,33], while some have additionally reported the motif classification [8,29]. We compiled a list of 18 CTD sequence motifs (Methods; **Table S8**) from the published literature and generated a motif similarity network (MSN) (**Figure S3**). From the MSN, we observed that the unclassified motifs clustered with the motifs classified as type A, whereas the motifs classified as type B are distinct (**Figure S3**). In addition, Kharade and McBride [11] reported a third type of CTD motif (type C) for the secreted proteins. Therefore, we accessed the HMM profiles from TIGRFAM [34] and NCBI protein family models (https://www.ncbi.nlm.nih.gov/protfam) for all three types of CTD (type A, type B and type C) to classify the compiled list of 102 secreted proteins. We observed that the HMM profiles and the motif-based classification of the 102 secreted proteins yielded similar results with 68 proteins as type A, 18 proteins as type B and 1 protein as type C (**Figure 4**). From the classified CTD sequences, we generated HMM profiles (**Supplemental Information**). Additionally, we report CTD motif sequences specific to the classified proteins (**Table 1**). Importantly, these motif sequences are better at classifying the different types of CTD in comparison with the existing sequence motifs (**Figure 4**).

**Figure 4:**
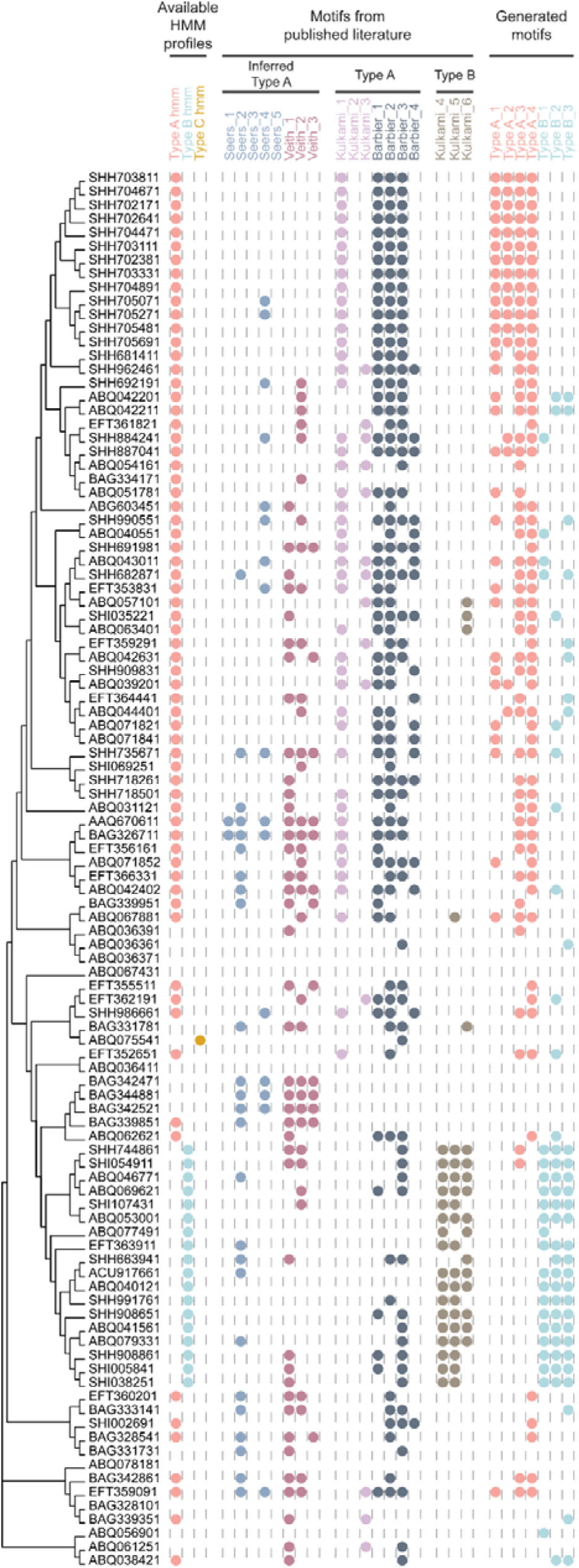
Identification and classification of the C-terminal domain (CTD; last 120 aa) of 102 experimentally characterized secreted proteins into three CTD types (type A, type B or type C) based on HMM profiles and sequence motifs from published literature. The motifs from Seers *et al.* [33] and Veith *et al.* [31] are classified as type A CTD based on the motif similarity network (**Figure S3**). Based on the classification of the secreted proteins, we generated 4 motifs for type A and 3 motifs for type B. The CTD of the secreted proteins are clustered using ClustalW [46] and the colored dots represent the presence of the CTD types. The figure was generated using Evolview [53] web server (https://www.evolgenius.info/evolview/).

**Table 1:** The table lists four type A and three type B CTD motifs generated in this study. We used the ‘discriminative mode’ in the MEME tool (https://meme-suite.org/meme/tools/meme) to generate the motifs from the input list of experimentally characterized secreted proteins. For each motif, we also provide the e-value given by the MEME tool.

| Motif Identifier | Motif | E-value |
| --- | --- | --- |
| Type A_1 | KIYPNPVSEILNIALQE[GN]LQL[EQ]KVN FYNTLGQLIKTTNHSE[IT]N<br>[VI]SSFAKG | 7.3e-454 |
| Type A_2 | VF[GR][ND]V[TN][QK]SNCALNVP[AT]GT[EQ]A[AV]YQAAAVW[KR]<br>][DN]FSPISG[SN]LLSNHSFAIES[NA]L | 2.6e-398 |
| Type A_3 | VK[VI]YPN[PQ]GKGT[LK][TN]I[SI] | 1.3e-158 |
| Type A_4 | [LV]SNLA[KS]G[VI]Y[IFL][VL]K[IV]XX | 7.4e-115 |
| Type B_1 | F[TS]PNGDGYNDT[WF]Y[IP]E[NG][IL]EXYP | 2.5e-091 |
| Type B_2 | ELP[SA]G[DT]Y[WY][YF][VI][LVI]K[YF]N[DE]N[KN][NT]XK | 6.4e-067 |
| Type B_3 | [VI]EI[FY][DN]R[YWF]G[KRV]L[IV][KY]ELDG[NYG]DNG[WD] | 6.1e-070 |

### Computational pipeline to predict proteins secreted via T9SS

We present the first computational pipeline to predict proteins secreted via T9SS across members of the *Bacteroidetes* phylum (**Figure 5**). The pipeline integrates several tools which predict important features in protein sequences. We used SignalP-5.0 [35] and Phobius [36] to detect the N-terminal Sec signal, SignalP-5.0 to detect N-terminal Tat signal, Phobius and TMHMM 2.0 [37] to detect the transmembrane domain (TM), and generated CTD HMM profiles to detect the CTD types in the protein sequences. We used ‘OR rule’ in the case where the feature is predicted by multiple tools. Based on the features of 102 experimentally identified proteins secreted via T9SS (**Table S3**), we designate a protein as secreted protein if: (i) it has Sec signal in N-terminal region; (ii) it has no Tat signal in N-terminal region; (iii) it has no TM domain; (iv) it has at least one of the three CTDs namely, type A, type B, or type C in its C-terminal region. We developed a computational tool implementing the above pipeline (**Figure 5**) to predict the secreted proteins from an input proteome. We used this tool to predict the secreted proteins for the 402 *Bacteroidetes* strains (**Table S5**) predicted to have T9SS. The predictions are available under the ‘Secreted protein prediction’ section of respective *Bacteroidetes* strain at: https://cb.imsc.res.in/t9gpred/.

**Figure 5:**
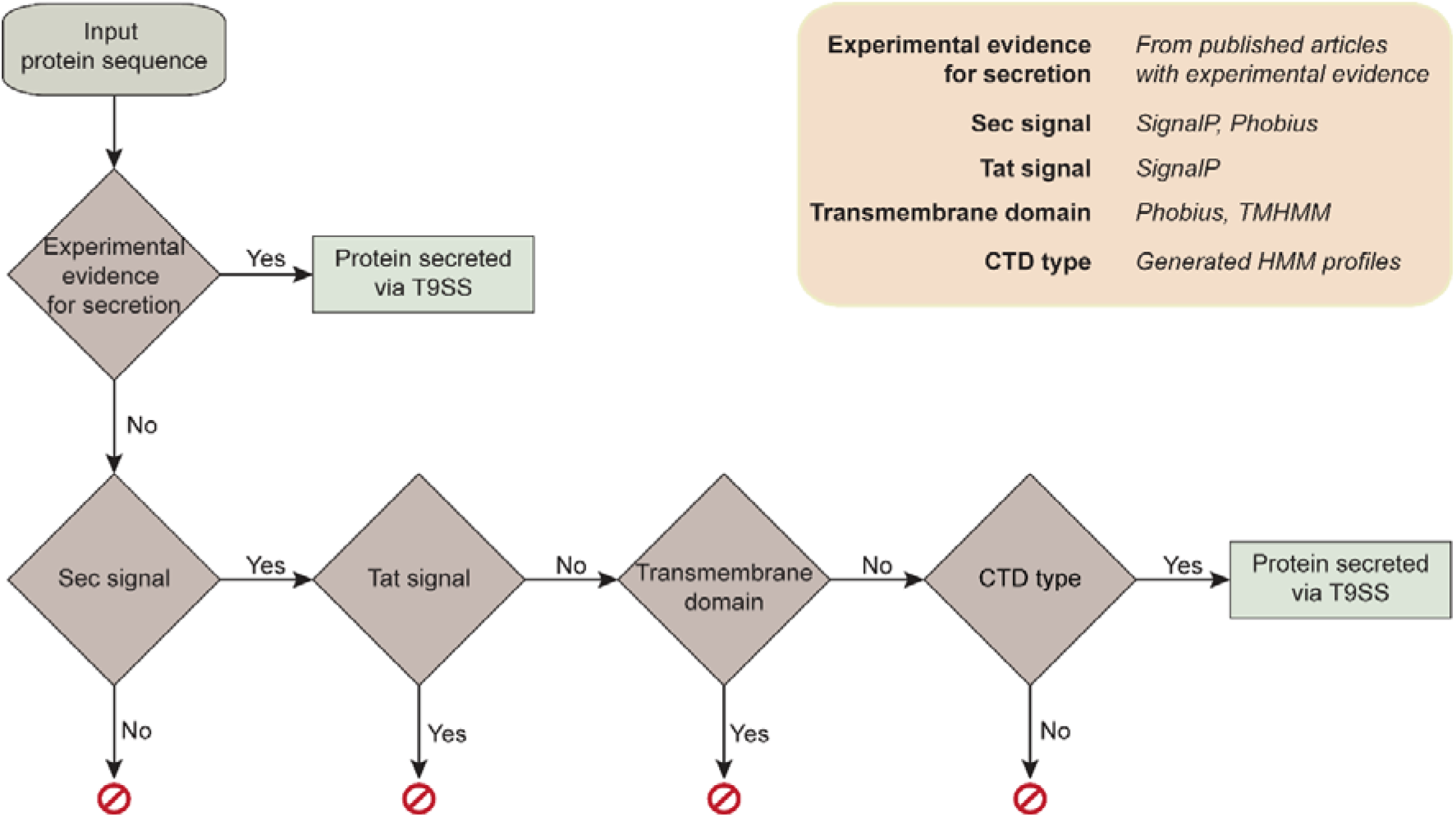
The computational pipeline to predict proteins secreted via T9SS. Computational tools used in this pipeline to predict features in the input protein sequence are mentioned in the top right panel in the figure.

### Validation of T9GPred predictions of secreted proteins

We validated the secreted protein prediction from T9GPred using recently published experimental evidence (after August 2020) on secreted proteins (**Supplemental Information**; **Figure S2**). We identified 22 new secreted proteins that were experimentally characterized from 4 different strains (**Table S9**). T9GPred predicted 14 of these 22 sequences to be secreted via T9SS. Among the 8 sequences that were not predicted by T9GPred, 2 lacked Sec signal sequences but contained type A CTD signal while the other 6 sequences did not have any of 3 CTD signals. Additionally, we estimated the precision of our CTD HMM prediction using 2 strategies (Methods). When checked against a total of 16,886 secreted proteins predicted by T9GPred, the first strategy always resulted in zero hits while the second strategy resulted in 972, 970, and 948 hits (average of 963 hits) yielding the precision of 100% and 94.29% respectively (Methods).

### Comparative analysis of the secreted proteins from gliding and non-gliding Bacteroidetes strains

Using our computational tool, we predicted 327 *Bacteroidetes* strains to have gliding motility (gliding *Bacteroidetes*) and 75 *Bacteroidetes* strains to lack gliding motility (non-gliding *Bacteroidetes*) (Methods). Further, we also predicted the secreted proteins for both gliding and non-gliding *Bacteroidetes* using our computational tool. From our predictions, we noted that gliding *Bacteroidetes* secrete more proteins than the non-gliding *Bacteroidetes*. To check for the statistical significance of this difference, we first normalized the number of secreted proteins with the total number of proteins (coding sequences) in the respective *Bacteroidetes* strains. We observed that gliding *Bacteroidetes* significantly secrete more proteins than non-gliding *Bacteroidetes* (*p*<0.001; corrected using Bonferroni method; **Figure 6a**). We also observed that the genome of gliding *Bacteroidetes* is significantly larger than non-gliding *Bacteroidetes* (*p*<0.01; corrected using Bonferroni method; **Figure 6b**).

**Figure 6:**
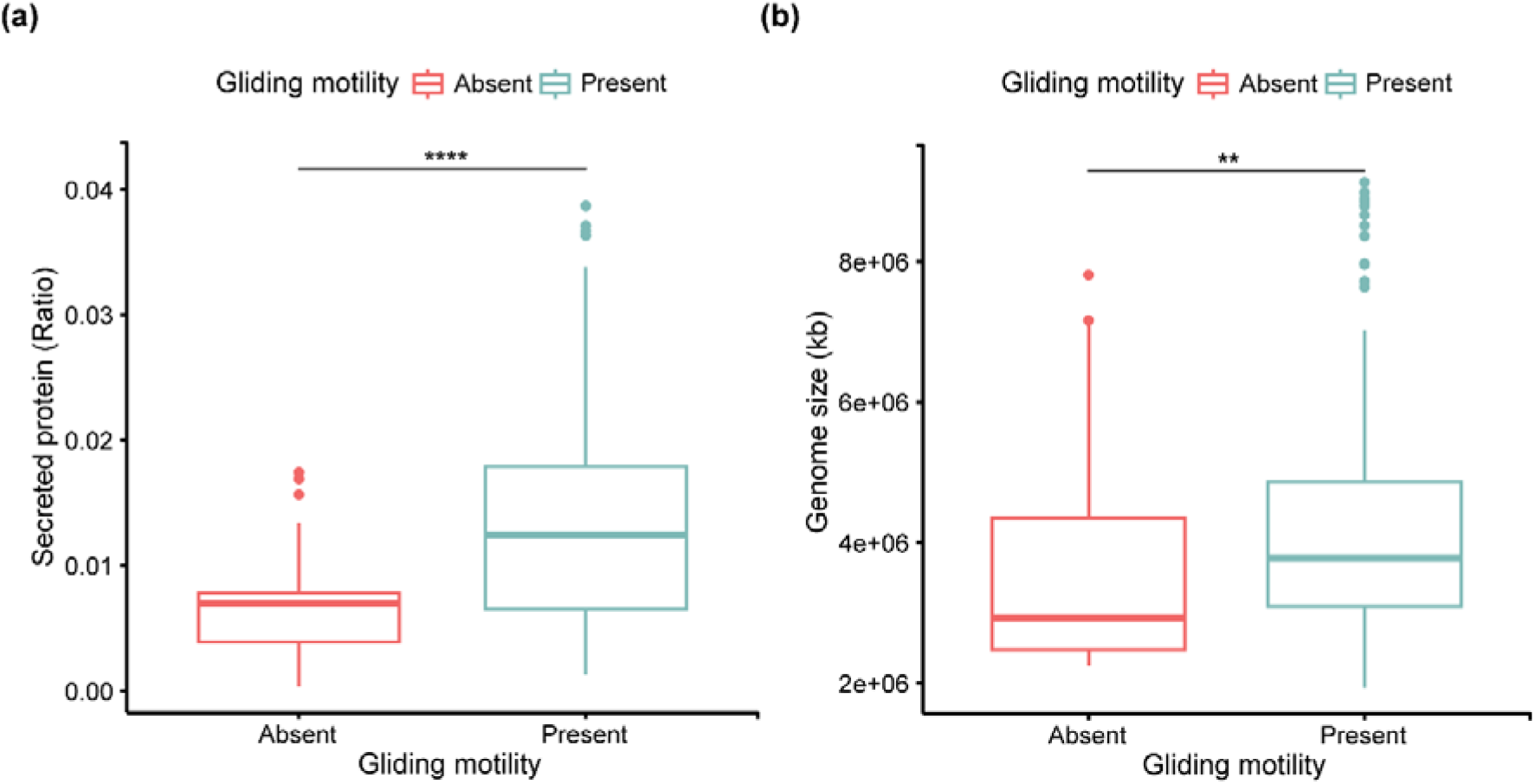
Comparative analysis of the secreted proteins and genome size between gliding and non-gliding *Bacteroidetes* strains predicted in this study. **(a)** Comparison of the predicted secreted proteins normalized with the total number of coding sequences of respective bacteria (ratio) between gliding and non-gliding *Bacteroidetes* strains (significantly different at *p*<0.001). **(b)** Comparison of the genome sizes (in kb) between gliding and non-gliding *Bacteroidetes* strains (significantly different at *p*<0.01). In each case, the significant difference between two samples is calculated using Bonferroni correction in rstatix package (https://cran.r-project.org/web/packages/rstatix/index.html) available in R version 4.2.1. Both plots are made using the ggplot2 [54] package in R version 4.2.1.

### T9GPred web server

We have designed a Findable, Accessible, Interoperable and Reusable (FAIR) [38] compliant user-friendly web server, **T**ype **9** secretion system and **G**liding motility **Pred**iction (T9GPred) that compiles the predictions and the computational tool developed in this study. The web server is available at: https://cb.imsc.res.in/t9gpred/. T9GPred is an organized web server with simple navigation that enables easy access to users (**Figure S4a**).

Users can interactively browse for the 693 *Bacteroidetes* strains in the ‘Browse’ section of the navigation bar. Users can hover over the heatmap to see information on the name of the bacteria, the predicted protein components and the number of predicted sequences of the protein component in the proteome of the bacteria (**Figure S4b**). Users can click on the name of the bacteria to see further details. The ‘Bacteroidetes details’ section gives the information on the organism name, genome identifier from PATRIC [34] or NCBI (https://www.ncbi.nlm.nih.gov/assembly/), isolation source of the bacteria, sequencing center information, predictions on T9SS, gliding motility and the number of secreted proteins, taxonomic information and proteome of the bacteria as a downloadable file (**Figure S4c**). Users can view the predicted protein components associated with T9SS on the ‘T9SS prediction’ page (**Figure S4d**) and the predicted protein components associated with gliding motility on the ‘Gliding motility prediction’ page (**Figure S4e**). For bacteria predicted to have T9SS, the predicted components can be visualized on the genome of the bacteria on the ‘Gene cluster visualization’ page (**Figure S4f**). Finally, the ‘Secreted protein prediction’ page lists the predicted secreted proteins and their annotations including the protein identifier from PATRIC (https://www.bv-brc.org/) or NCBI (https://www.ncbi.nlm.nih.gov/protein/), UniProt [39] identifier, AlphaFold DB [40,41] identifier, type of evidence, type of CTD and the presence of Carbohydrate-Active enZYmes (CAZy) domain predicted using dbCAN2 tool [42] (**Figure S4g**).

Users can access the prediction tools in the ‘Prediction’ section of the navigation bar. The ‘Prediction of T9SS and gliding motility’ page contains the tool for the prediction of T9SS or gliding motility in the input proteome (**Figure S5a**). Users can upload the proteome in fasta file format or paste the sequence in the provided text box to run the prediction tool. The tool returns the predicted protein components along with the decision for the presence of T9SS or gliding motility (**Figure S5a**). The ‘Prediction of protein secreted via T9SS’ page allows users to predict whether the input protein can be secreted via T9SS (**Figure S5b**). Users can paste the sequence in the provided text box to run the prediction tool. The tool returns the prediction of different features including the classification of CTD types and mentions whether the protein can be secreted via T9SS (**Figure S5b**).

## DISCUSSION

T9SS is one of the least characterized secretion systems in Gram-negative bacteria (**Supplemental Information**; **Figure S6**). In this study, we leveraged the existing experimental evidence to develop the first comprehensive computational tool, T9GPred, which predicts the presence of T9SS, gliding motility and proteins secreted via T9SS from a bacterial proteome. In particular, we designated 6 key proteins (GldK, GldL, GldM, GldN, SprA and SprE) as mandatory for a functional T9SS and observed that they are sufficient for T9SS prediction when checked against the proteome of recently characterized *Bacteroidetes* strains having T9SS (**Table S6**). Using these 6 T9SS mandatory proteins, we predicted 402 of the 693 completely sequenced *Bacteroidetes* strains to have T9SS. Additionally, we also observed that our tool T9GPred predicted the presence of T9SS only in the *Bacteroidetes* phylum (**Supplemental Information**; **Table S10**) which is in line with the findings by McBride and colleagues [1,3]. Further, we also observed that the T9SS mandatory protein components are clustered in the bacterial genome (**Supplemental Information**; **Figure S7**) suggesting that GldK, GldL, GldM and GldN form an operon in members of the *Bacteroidetes* phylum having T9SS [12,24].

In some *Bacteroidetes* strains, T9SS also aids in the movement of the species which is termed as gliding motility [1,3]. In addition to the 6 T9SS mandatory proteins, we designated 5 key proteins (GldB, GldD, GldH, GldJ and SprT) as mandatory for exhibiting gliding motility and observed that they are sufficient for gliding motility prediction when checked against the proteome of recently characterized *Bacteroidetes* strains exhibiting gliding motility (**Table S7**). Using these 11 gliding motility mandatory proteins, we predicted 327 of 402 *Bacteroidetes* strains with T9SS to also have gliding motility. Moreover, we observed that these 327 *Bacteroidetes* strains do not have flagellar motor or type IV pili (**Supplemental Information**). We also noted that there is no overlap between the protein components associated with gliding motility and flagellar motor or type IV pili (**Supplemental Information**). Thus, our analysis suggests that gliding motility associated with T9SS is unique to the *Bacteroidetes* phylum.

The proteins secreted via T9SS are involved in multiple processes including the survival and pathogenicity of the bacteria [13,28]. We identified 3 types of C-terminal domain (CTD) signals (type A, type B or type C) by leveraging the reported signal sequences [8,29,31,33] in conjunction with published evidence. We leveraged this CTD classification to build a computational pipeline for secreted protein prediction and found its predictions are highly precise and agree well with recently published experimental evidence (**Table S9**). A comparative analysis with the proteins secreted via other *Bacteroidetes*-specific secretion systems highlighted the uniqueness of the CTD signal in the proteins secreted via T9SS (**Supplemental Information**). We found that members of the *Bacteroidetes* phylum exhibiting gliding motility have significantly larger genome sizes and secrete more proteins in comparison with the members that do not have gliding motility (**Figure 6**). This suggests that members of the *Bacteroidetes* phylum exhibiting gliding motility have a larger genome probably encoding more specialized proteins to aid in gliding motility.

T9GPred has its fair share of limitations like any other computational tool. T9GPred does not mention the accuracy of the predictions as it lacks information on negative control. It relies on protein profiles trained on a small set of experimentally characterized *Bacteroidetes* strains to date, which limits its ability to exhaustively capture all variations. Moreover, T9GPred does not comment on the maturation of the secreted proteins as it does not include the information on the functional characterization of accessory protein components. The secreted protein prediction by T9GPred is limited by the choice of 3 CTD types considered in this study which is not indicative of all possible secreted proteins [8]. Moreover, T9GPred does not include the structural conservation of the CTD [43] but considers only the protein sequence for the prediction of the secreted proteins. T9GPred predicts putative secreted proteins but does not comment on the variation in the levels of secreted proteins in different environments.

Nevertheless, T9GPred aids in streamlining experiments by providing quick predictions. For pathogens, the predicted T9SS protein components and the secreted proteins could be targeted for drug discovery. For non-pathogens, T9GPred predicts secreted proteins that could be the putative subjects for genetic enhancements. Importantly, T9GPred predictions are robust to the draft genome sequences. T9GPred has the ability to identify a new uncharacterized bacterium as belonging to the *Bacteroidetes* phylum. In conclusion, we developed T9GPred, a computational tool that not only predicts T9SS but also predicts T9SS based gliding motility and the proteins secreted via T9SS. We have made the predictions and the tool available on our web server which is accessible at: https://cb.imsc.res.in/t9gpred/. The source code of the computational tool is available at: https://github.com/asamallab/T9GPred.

## METHODS

### Workflow to identify proteins associated with T9SS or gliding motility among members of the *Bacteroidetes* phylum

We mined PubMed (https://pubmed.ncbi.nlm.nih.gov/) to compile an exhaustive list of experimentally characterized proteins associated with T9SS or gliding motility in members of the *Bacteroidetes* phylum from published literature. We queried PubMed using the following keywords:

(“type 9 secretion system”[Title/Abstract] OR “type 9 secretion systems”[Title/Abstract] OR “type IX secretion system”[Title/Abstract] OR “type IX secretion systems”[Title/Abstract] OR “type-9 secretion system”[Title/Abstract] OR “type-9 secretion systems”[Title/Abstract] OR “type-IX secretion system”[Title/Abstract] OR “type-IX secretion systems”[Title/Abstract] OR “t9ss”[Title/Abstract] OR “T9SS”[Title/Abstract] OR “PoRSS”[Title/Abstract] OR “PoR secretion system”[Title/Abstract] OR “Type 9 secretion”[Title/Abstract] OR “Type IX secretion”[Title/Abstract])

and retrieved 107 published articles on 26 August 2020 (**Figure 1**). We noted that our PubMed query was unable to retrieve some previous studies focussing on gliding motility and the type 9 based secretion, as such studies were published prior to the use of T9SS nomenclature. Therefore, we manually compiled 22 additional published articles on gliding motility or type 9 based secretion (**Figure 1**). Thereafter, we excluded the articles which are computational studies or reviews; or which did not contain mutation-based experiments. Finally, we shortlisted 47 published articles from which we identified 23 proteins associated with T9SS (**Table S1**), 22 proteins associated with gliding motility (**Table S2**) and 102 proteins secreted via T9SS (secreted proteins) in members of the *Bacteroidetes* phylum (**Table S3**). **Figure 1** summarizes the workflow according to the PRISMA statement [44].

### Compilation of completely sequenced *Bacteroidetes* strains

Our aim is to predict the presence of T9SS or gliding motility across the *Bacteroidetes* phylum. Thus, we considered *Bacteroidetes* strains for which the complete genome is available. We queried NCBI assembly (https://www.ncbi.nlm.nih.gov/assembly) using the following keywords:

(“Bacteroidetes”[Organism] or “Bacteroidia”[Organism] or Bacteroidetes[All Fields]) and (bacteria[filter] and “latest refseq”[filter] and “complete genome”[filter] and all[filter] not anomalous[filter])

and identified 693 completely sequenced *Bacteroidetes* strains on 21 December 2020 (**Table S5**). For these 693 *Bacteroidetes* strains, we accessed the corresponding proteome from PATRIC [45] (https://www.patricbrc.org/) or NCBI assembly (https://www.ncbi.nlm.nih.gov/assembly). Note that if the proteome for a bacterium was available in both PATRIC and NCBI assembly, we preferred the proteome from PATRIC.

### Bioinformatics analysis to identify and predict T9SS or gliding motility

Among the 693 completely sequenced *Bacteroidetes* strains, we identified 43 *Bacteroidetes* strains to have experimentally or computationally characterized proteins associated with T9SS or gliding motility from published literature (**Table S11**; **Supplemental Information**). Further, we used these 43 proteomes to generate Hidden Markov Model (HMM) profiles for the identified protein components associated with T9SS or gliding motility. The detailed workflow to generate HMM profiles is documented in the **Supplemental Information**. Subsequently, we relied solely on the experimental evidence from 30 *Bacteroidetes* strains to infer the mandatory protein components for T9SS and gliding motility (**Table S4**). We collated the HMM profiles of the designated mandatory components and scripted a python program that checks for the presence of T9SS or gliding motility across any sequenced *Bacteroidetes* strains.

### Computational analysis of secreted proteins

We collected the sequences of the experimentally identified 102 secreted proteins (**Table S3**) from NCBI GenBank (https://www.ncbi.nlm.nih.gov/genbank/). We considered the last 120 amino acids of each protein sequence as their C-terminal domain (CTD) [8]. Further, we used ClustalW [46] with default parameters to align the CTD sequences of the 102 secreted proteins. Thereafter, we clustered the aligned sequences using MEGA X [47] with the maximum-likelihood method and default parameters.

We relied on HMM profile based prediction and motif analysis to classify the 102 CTD sequences. We accessed the HMM profiles for CTD types from TIGRFAM [34] and NCBI protein family models (https://www.ncbi.nlm.nih.gov/protfam) and compiled 18 CTD sequence motifs of secreted proteins from published literature (**Table S8**). To find the similarity between the motifs, we created a motif similarity network (MSN) using Tomtom [48] (https://meme-suite.org/meme/tools/tomtom) by setting the overlap >= 3 and score >= 2 [49] and visualized the MSN using Cytoscape [50] (**Figure S3**). Further, we determined the presence of the motifs in the CTD of the secreted proteins using Find Individual Motif Occurrence (FIMO) (https://meme-suite.org/meme/tools/fimo) tool [51] with default parameters. We classified the CTD types of secreted proteins based on the HMM prediction and substantiated with motif based classification. Using the classified CTD sequences, we created the corresponding HMM profiles (**Supplemental Information**) and generated motifs using Multiple Expectation Maximization for Motif Elicitation (MEME) [52] (https://meme-suite.org/meme/tools/meme) tool in the discriminative mode of motif discovery.

### Checking for random hits of CTD HMM profiles

We followed Veith *et al.* [31] to check for random hits of the 3 CTD HMM profiles (type A, type B and type C) on proteome of the 402 *Bacteroidetes* strains using 2 different strategies: (i) randomizing the CTD regions (last 120 amino acids) of each protein while maintaining amino acid composition; (ii) randomly selecting 120 amino acid long sequence from the full sequence of each protein. Since we considered the CTD sequence length as 120 amino acids, we filtered out those predicted protein sequences that were less than or equal to 120 amino acids to be consistent with the randomization strategy. We generated three data sets for each strategy to estimate the number of random hits. We then used the following formula to calculate the precision: precision -

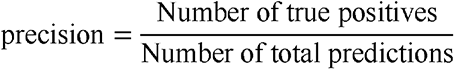

where we estimated the Number of true positives as:

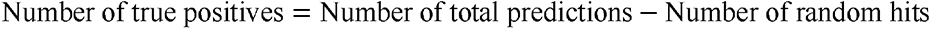

## Web interface and web server management system

We created a user-friendly web server **T**ype **9** secretion system and **G**liding motility **Pred**iction (T9GPred) to compile and share the predictions and the computational tool generated in this study. T9GPred compiles the proteins associated with T9SS or gliding motility in the 693 completely sequenced *Bacteroidetes* strains. For the *Bacteroidetes* strains predicted to have T9SS, T9GPred lists the proteins predicted to be secreted via T9SS. Moreover, T9GPred provides a prediction tool to check for the presence of T9SS or gliding motility, and secreted proteins in any sequenced genome. T9GPred is openly available at https://cb.imsc.res.in/t9gpred/.

To create the web server, we used MariaDB (https://mariadb.org/) to store the compiled information and Structured Query Language (SQL) to retrieve information. We used PHP (https://www.php.net/) with custom HTML, CSS, jQuery (https://jquery.com/), Bootstrap 4 (https://getbootstrap.com/docs/4.0/), and Plotly JavaScript (https://plotly.com/javascript/) to create the web interface of T9GPred. T9GPred web server is hosted on an Apache (https://httpd.apache.org/) server running on Debian 9.4 Linux Operating System.

## DATA AND CODE AVAILABILITY

The predictions and the computational tool are accessible via the web server, T9GPred available at: https://cb.imsc.res.in/t9gpred/. The standalone version of the developed computational tool and the codes to generate different plots are available in our GitHub repository: https://github.com/asamallab/T9GPred.

## AUTHOR CONTRIBUTIONS

**Ajaya Kumar Sahoo:** Conceptualization, Data Compilation, Data Curation, Formal Analysis, Software, Visualization, Writing; **R. P. Vivek-Ananth:** Conceptualization, Formal Analysis, Software, Visualization, Writing; **Nikhil Chivukula:** Conceptualization, Formal Analysis, Software, Visualization, Writing; **Shri Vishalini Rajaram:** Conceptualization, Data Compilation, Data Curation, Formal Analysis; **Karthikeyan Mohanraj:** Software, Visualization; **Devanshi Khare:** Formal Analysis; **Celin Acharya:** Conceptualization, Formal Analysis, Writing; **Areejit Samal:** Conceptualization, Supervision, Formal Analysis, Software, Writing.

## ACKNOWLEDGEMENTS

We thank Gokul Balaji Dhanakoti and B. Raveendra Reddy for computational support, and Kishan Kumar for help in generating a figure. We thank Anshu Bhardwaj, Dhiraj Kumar, Mitali Mukerji, Vinay Nandicoori, Hema Rajaram and Smita Srivastava for discussions. Areejit Samal would like to acknowledge support from the Department of Atomic Energy (DAE), Government of India (GoI), the Science and Engineering Research Board (SERB), GoI [Ramanujan Fellowship SB/S2/RJN-006/2014], and the Max Planck Society, Germany [Max Planck Partner Group in Mathematical Biology]. The funders have no role in the study design, data collection, data analysis, manuscript preparation, or decision to publish.

## Supplemental Information

Supplemental Information consists of Supplemental Text, Supplemental Figures S1-S8 and Supplemental Tables S1-S13.

### Supplemental Figure Captions

**Figure S1: (a)** Distribution of the number of proteins associated with T9SS in each of 402 *Bacteroidetes* strains predicted to have T9SS. The mean and median of the distribution are 19.30 and 19, respectively. **(b)** Distribution of the number of proteins associated with T9SS or gliding motility in each of 327 *Bacteroidetes* strains predicted to exhibit gliding motility. The mean and median of the displayed distribution are 21.98 and 22, respectively.

**Figure S2:** Workflow to compile new experimental evidence on T9SS, gliding motility and associated secreted proteins. This workflow is presented according to the PRISMA statement.

**Figure S3:** Motif similarity network (MSN) of the 18 compiled CTD motifs generated using Tomtom software by setting the overlap >= 3 and score >= 2. We visualized the MSN using Cytoscape, where each node represents the motifs and the edges represent the score computed using Tomtom.

**Figure S4:** Snapshots of the web interface of the T9GPred web server. (a) ‘Home’ page showing navigation bar. (b) ‘Browse’ option enables users to explore the details and predictions for 693 completely sequenced *Bacteroidetes* strains considered in this study. (c) The detailed information page for selected *Bacteroidetes* strain. (d) The ‘T9SS prediction’ page lists the predicted protein components associated with T9SS and visualizes the predicted components. (e) The ‘Gliding motility prediction’ page lists the predicted protein components associated with gliding motility and visualizes the predicted components. (f) The ‘Gene cluster visualization’ page provides a visualization of the location of the predicted protein components in the genome of the selected *Bacteroidetes strain*. (g) The ‘Secreted protein prediction’ page tabulates the proteins predicted to be secreted via T9SS of the selected *Bacteroidetes* strain.

**Figure S5:** Snapshots of the prediction tool in T9GPred web server. (a) The ‘Prediction of T9SS and gliding motility’ page allows users to submit the proteome of a bacterium and provides the prediction on the T9SS and gliding motility in a tabular format. (b) The ‘Prediction of protein secreted via T9SS’ page allows users to submit a protein sequence and provides the prediction of the features in a tabular format.

**Figure S6:** Cumulative estimate of published research articles for the nine different secretion systems (T1SS-T9SS) in bacteria. We have not considered T10SS or T11SS in this estimate as these two systems are very recently discovered.

**Figure S7:** Gene cluster visualization of the proteins associated with T9SS or gliding motility in the genome of *Flavobacterium johnsoniae* UW101 and *Porphyromonas gingivalis* ATCC 33277. The operons consisting of GldK (1) GldL (2) GldM (3) GldN (4) are shown in the enlarged view. The figure was generated using CGView package.

**Figure S8:** Flowchart describing the workflow to create the HMM profile from protein sequences.

### Supplemental Tables

**Table S1:** The table gives the compiled list of 23 proteins with experimental evidence of being associated with T9SS in members of the *Bacteroidetes* phylum. For each protein, the table gives the GenBank protein accession from respective *Bacteroidetes* strain, type of experimental evidence, and PubMed identifier of literature reference(s).

**Table S2:** The table gives the compiled list of 22 proteins with experimental evidence of being associated with gliding motility in members of the *Bacteroidetes* phylum. For each protein, the table gives the GenBank protein accession from respective *Bacteroidetes* strain, type of experimental evidence, and PubMed identifier of literature reference(s).

**Table S3:** The table gives the compiled list of 102 proteins with published experimental evidence of being secreted via T9SS in members of *Bacteroidetes* phylum. For each protein, the table gives the GenBank protein accession, *Bacteroidetes* strain in which the protein was identified, type of experimental evidence, and PubMed identifier of literature reference(s). Moreover, the table gives for each protein sequence, the Sec signal prediction from SignalP and Phobius, Tat signal prediction from SignalP, and the Transmembrane domain prediction from Phobius and TMHMM.

**Table S4:** The table gives the list of 30 *Bacteroidetes* strains with experimental evidence on the presence or absence of T9SS or gliding motility. For each *Bacteroidetes* strain, the table also gives PubMed identifier of literature reference(s).

**Table S5:** The table gives the compiled list of 693 completely sequenced *Bacteroidetes* strains considered for this study. For each *Bacteroidetes* strain, the table provides taxonomic information, isolation source, sequencing center information, prediction of T9SS, gliding motility and the number of proteins predicted to be secreted via T9SS.

**Table S6:** The table gives the compiled list of new *Bacteroidetes* strains having experimental evidence for the presence of T9SS in their genome. For each strain, the table gives the genome identifier, genome assembly level, T9GPred predictions and PubMed identifier of the literature reference. Note, T9GPred also predicted all the 4 strains to have T9SS and 2 of 4 strains are of new species which were not considered while building T9GPred.

**Table S7:** The table gives the compiled list of new *Bacteroidetes* strains having experimental evidence for the presence of gliding motility in their genome. For each strain, the table gives the genome identifier, genome assembly level, T9GPred prediction and PubMed identifier of the literature reference. Note, T9GPred also predicted all the 7 strains to have gliding motility and 4 of 7 strains are of new species which were not considered while building T9GPred.

**Table S8:** The table gives the list of 18 sequence motifs for the C-terminal domain (CTD) of the proteins secreted via T9SS. For each motif, the table gives the motif identifier which is based on the author name, type of CTD motif and the PubMed identifier of the literature reference.

**Table S9:** The table gives the compiled list of 22 new secreted proteins that were experimentally characterized from 4 different strains. For each protein, the table gives the respective *Bacteroidetes* strain name, experimental evidence, T9GPred prediction for secreted protein and PubMed identifier of the literature reference.

**Table S10:** The table gives the list of 34,369 completely sequenced bacteria from NCBI assembly. For each bacterium, the table gives the phylum information and the prediction of T9SS using T9GPred. Note that the phylum *Bacteroidetes* and *Bacteroidota* are synonyms according to NCBI Taxonomy 2023 (https://ncbiinsights.ncbi.nlm.nih.gov/2022/11/14/prokaryotic-phylum-name-changes/).

**Table S11:** The table gives the compiled list of 43 *Bacteroidetes* strains considered to generate Hidden Markov Model (HMM) profiles for 28 proteins associated with T9SS or gliding motility. For each *Bacteroidetes* strain, the table provides the genome identifier, true positive hit(s) of the 28 proteins, type of evidence of protein characterization and the PubMed identifier of literature reference(s).

**Table S12:** The table gives the PubMed query for the each of the nine different secretion systems (T1SS-T9SS) in bacteria and the respective number of articles retrieved. The PubMed search was last performed on 10 January 2021.

**Table S13:** The table gives the true positive hit for the type C C-terminal domain (CTD) generated from analyzing 693 completely sequenced *Bacteroidetes* strains.

