## Supplemental Text and Figures S1-S8 for "T9GPred: A Comprehensive Computational Tool for the Prediction of Type 9 Secretion System, Gliding Motility and the Associated Secreted Proteins"

### for

### Supplemental Text

#### Type 9 secretion system (T9SS) is one of the least characterized secretion systems in Gram-negative bacteria

T9SS is one of the recently discovered secretion systems in Gram-negative bacteria [1–3]. There are ten other secretion systems of which T10SS and T11SS are very recently discovered and less understood [4–7]. To gauge the level of existing information on the remaining nine secretion systems (T1SS-T9SS), we estimated the number of published articles for each case using a similar PubMed (https://pubmed.ncbi.nlm.nih.gov/) query. **Table S12** lists the query and the number of published articles in PubMed for each of the nine secretion systems. It is apparent that T9SS is a recently discovered secretion system with fewer published articles in comparison to other secretion systems (except T8SS) (**Figure S6**).

Though TXSScan [8,9] can predict presence of T9SS (among other secretion systems), it does not predict the presence of gliding motility or the proteins secreted via T9SS. In this study we identified protein components associated with T9SS and gliding motility which are not captured by TXSScan. Our efforts led to the development of computational tool which can predict T9SS, gliding motility and secreted proteins among members of the *Bacteroidetes* phylum.

#### Workflow to create Hidden Markov Model profiles

Hidden Markov Model (HMM) profiles of protein sequences are used to computationally predict the presence of the protein in the proteome of the organism where such protein is not previously known to be present [10–12]. In this study, we generated HMM profiles for: (i) proteins associated with T9SS or gliding motility; (ii) different C-terminal domain (CTD) types of secreted proteins. The steps involved in generating a HMM profile are as follows:

1. Identify protein for which the HMM profile is to be generated
2. Multiple sequence alignment (MSA) of all the sequences of an identified protein using the MAFFT tool [13]
3. Trimming the ends of MSA using Jalview software [14] to obtain the most conserved region
4. Generating HMM profile of the trimmed MSA using the HMMER tool [15] (http://hmmer.org/)
5. HMM search using the generated HMM profile against target proteome
6. Consider top results from the HMM search until one encounters a sequence that is *not* characterized as 'true positive'
7. **Repeat** steps 2-6 with the top results as the input sequence until the top results include all the true positives
8. Assign a GA (gathering) score [16] for the HMM profile such that all true positives are included

Note that for identifying the 'true positives', we used reciprocal BLAST [17] between the protein of interest and target proteome. The flowchart for generating the HMM profile is given in **Figure S8**.

*HMM profile generation for proteins associated with T9SS or gliding motility*

We compiled a list of proteins associated with T9SS or gliding motility from published literature. For each of the compiled proteins, we collected the experimentally characterized sequences. We considered such sequences as the input sequences for HMM profile generation. Further, we considered the target proteome from 43 of the 693 completely sequenced *Bacteroidetes* strains which are either experimentally or computationally characterized (**Table S11**).

In order to generate true positives, we performed BLAST [17] in two ways: (i) experimentally characterized sequences as input against the proteome of the 43 *Bacteroidetes* strains; (ii) proteome of 43 *Bacteroidetes* strains as input sequences against the experimentally characterized proteins; using BLASTp (https://blast.ncbi.nlm.nih.gov/Blast.cgi? PAGE=Proteins). In both ways, the e-value cut-off in BLAST result was considered to be the same. The considered e-values were based on previously reported studies and on the domain conservation between input and resultant protein sequences from Conserved Domains Database (CDD) [18]. When the experimentally characterized protein is mapped to the corresponding protein(s) from the proteome in both the BLAST results, we consider the mapping to satisfy the reciprocal BLAST. We manually verified all the proteins of the proteome that satisfy the reciprocal BLAST with the published studies, results from TIGRFAM HMM [12] search and genome annotations in PATRIC [19] and NCBI assembly (https://www.ncbi.nlm.nih.gov/assembly/). We removed sequences that could not be validated and added additional protein sequences that are reported to be homologous but were not identified through reciprocal BLAST due to poor conservation. This constitutes the true positives for a given protein. The true positives of the experimentally characterized proteins for the 43 *Bacteroidetes* strains are given in **Table S11**.

We followed the steps 1-8 of *Workflow to create Hidden Markov Model profiles* to generate the HMM profiles of the experimentally characterized proteins.

*HMM profile generation for CTD types of secreted proteins*

We classified the CTD of the 102 experimentally characterized secreted proteins into three CTD types namely, type A, type B and type C (Methods). For type A and type B, we followed steps 2-4 of *Workflow to create Hidden Markov Model profiles* to generate the HMM profiles from the classified CTD sequences. Since the generated HMM profiles for type A and type B did not contain GA score, we used an e-value cutoff of 1e-06 to perform HMM search.

For type C, we observed that there was only one classified protein (ChiA) among the 102 experimentally characterized secreted proteins (**Figure 4**). Initially, we relied on the 43 *Bacteroidetes* strains but could not find any protein homologous to ChiA. Therefore, we considered the 693 completely sequenced *Bacteroidetes* strains as the target proteome and generated the true positives for type C using the methodology described in *HMM profile generation for proteins associated with T9SS or gliding motility*. Thereafter, we generated type C HMM profile by following steps 1-8 of *Workflow to create Hidden Markov Model profiles*. The true positives of the type C proteins across *Bacteroidetes* strains are given in **Table S13**.

**T9SS is found only in the *Bacteroidetes* phylum**

McBride and colleagues had earlier observed that T9SS is unique to only the *Bacteroidetes* phylum and is not found in other phyla [1,3]. But there is no published extensive analysis where a check was made for the presence of T9SS across all the sequenced bacteria. Therefore, we were interested in using our computational tool to check for the presence of T9SS in sequenced bacteria. We queried NCBI Assembly (https://www.ncbi.nlm.nih.gov/assembly/) using the keywords:

("Bacteria"[Organism] OR bacteria[All Fields]) AND (latest[filter] AND "complete genome"[filter] AND all[filter] NOT anomalous[filter])

and retrieved 34,369 completely sequenced genomes (**Table S10**) on 28 February 2023. After that, we checked for the presence of the 6 designated T9SS mandatory components and observed that only the bacteria belonging to the *Bacteroidetes* phylum have all 6 components (**Table S10**).

### Four T9SS mandatory proteins form an operon-like cluster

We visualized the location of the predicted protein components associated with T9SS or gliding motility in their respective bacterial genome. We observed that 4 of the 6 T9SS mandatory proteins (GldK, GldL, GldM, GldN) are clustered together in the genome of *Bacteroidetes* strains predicted to have T9SS. Earlier studies have shown that GldK, GldL, GldM and GldN proteins form an operon in the genome of *Flavobacterium johnsoniae* [20] and *Porphyromonas gingivalis* [21]. This suggests that these 4 proteins may form an operon in other members of the *Bacteroidetes* phylum. **Figure S7** shows the visualization of the protein location in the genome of *F. johnsoniae* and *P. gingivalis* generated using CGView package [22,23]. Similar visualizations are available in the T9GPred web server, under the 'Gene cluster visualization' section of respective *Bacteroidetes* strains.

### *Bacteroidetes* strains having gliding motility lack other kinds of motility machinery

We checked for the presence of other motility machinery in the 327 *Bacteroidetes* strains predicted to have gliding motility by T9GPred. We used the HMM profiles for flagellar motor, type IVa pili (T4aP) and type IVb pili (T4bP) from TXSScan [8,9], and observed that none of the 327 *Bacteroidetes* strains have either flagellar motors or type IV pili. Further, we found that there was no overlap between the compiled list of 22 experimentally characterized proteins associated with gliding motility and the flagellar motor or type IV pili proteins. This suggests that *Bacteroidetes* strains having gliding motility do not have other motility machinery. The proteins associated with gliding motility provide unique machinery to aid movement in members of the *Bacteroidetes* phylum.

### Proteins secreted via T9SS have a unique secretion signal

Members of the *Bacteroidetes* phylum are reported to have other secretion systems apart from T9SS, namely T1SS, T4SS, T5SS and T6SS [8,24–26]. In this context, we probed the uniqueness of proteins secreted via T9SS in comparison with the secreted proteins from other secretion systems. We leveraged the generated HMM profiles of the T9SS CTD signals (type A, type B and type C) to check for their presence in the proteins secreted via the other secretion systems. We accessed: (i) 1024 proteins secreted via T1SS from Linhartová *et al.* [27]; (ii) 540 proteins secreted via T4SS from Bi *et al.* [28]; (iii) 47 proteins secreted via T5SS from Celik *et al.* [29]; (iv) 294 proteins secreted via T6SS from Li *et al.* [30]. We observed that the T9SS specific CTD signals are absent in the proteins secreted via other secretion systems which suggest that the CTD signal is unique to proteins secreted via T9SS.

**Curation and compilation of new experimental evidences**

To validate the T9GPred predictions, we relied on the experimental evidence on T9SS, gliding motility and associated secreted proteins, which were published after August 2020 (i.e., after the last date on which query was made to build T9GPred). We queried PubMed (https://pubmed.ncbi.nlm.nih.gov/) for experimental evidence on T9SS and associated secreted proteins using the following keywords:

("2020/06/01"[Date - Publication] : "3000"[Date - Publication]) AND ('type 9 secretion system*'[TIAB] OR 'Type 9 secretion system*'[TIAB] OR 'T9SS'[TIAB] OR 't9ss'[TIAB] OR 'type IX secretion*'[TIAB])

and retrieved 67 new articles on 27 April 2023 (**Figure S2**). We also queried PubMed for experimental evidence on gliding motility using the following keywords:

("2020/06/01"[Date - Publication] : "3000"[Date - Publication]) AND "gliding motility"

and retrieved 110 new articles on 27 April 2023 (**Figure S2**). We manually screened these new articles for experimental characterization of T9SS and associated secreted proteins or microscopic observation for gliding motility. We shortlisted 12 articles containing comprehensive experimental evidence (**Figure S2**). From the shortlisted articles, we identified 4 new *Bacteroidetes* strains to have T9SS (**Table S6**), 7 new *Bacteroidetes* strains to have gliding motility (**Table S7**) and 22 new proteins secreted via T9SS (**Table S9**). Note that some of the recently characterized *Bacteroidetes* strains have a draft genome sequence.

### Supplemental Figures

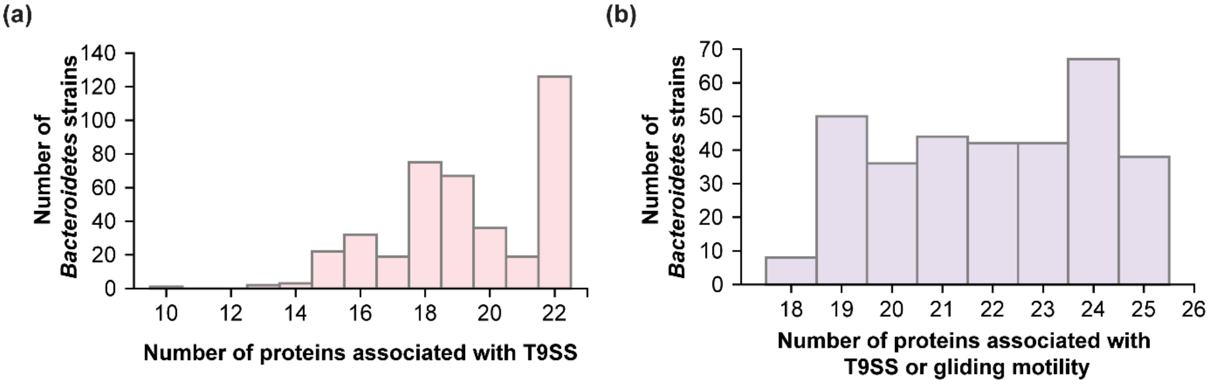


**Figure S1: (a)** Distribution of the number of proteins associated with T9SS in each of 402 *Bacteroidetes* strains predicted to have T9SS. The mean and median of the distribution are 19.30 and 19, respectively. **(b)** Distribution of the number of proteins associated with T9SS or gliding motility in each of 327 *Bacteroidetes* strains predicted to exhibit gliding motility. The mean and median of the displayed distribution are 21.98 and 22, respectively.

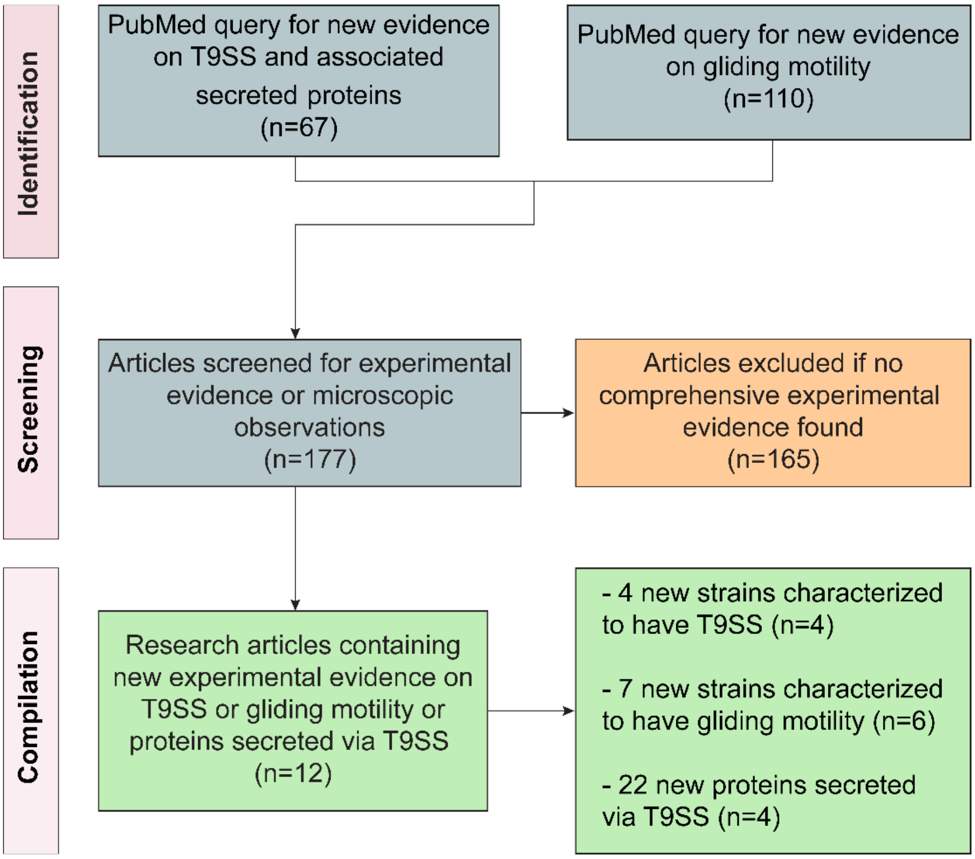


**Figure S2:** Workflow to compile new experimental evidence on T9SS, gliding motility and associated secreted proteins. This workflow is presented according to the PRISMA statement.

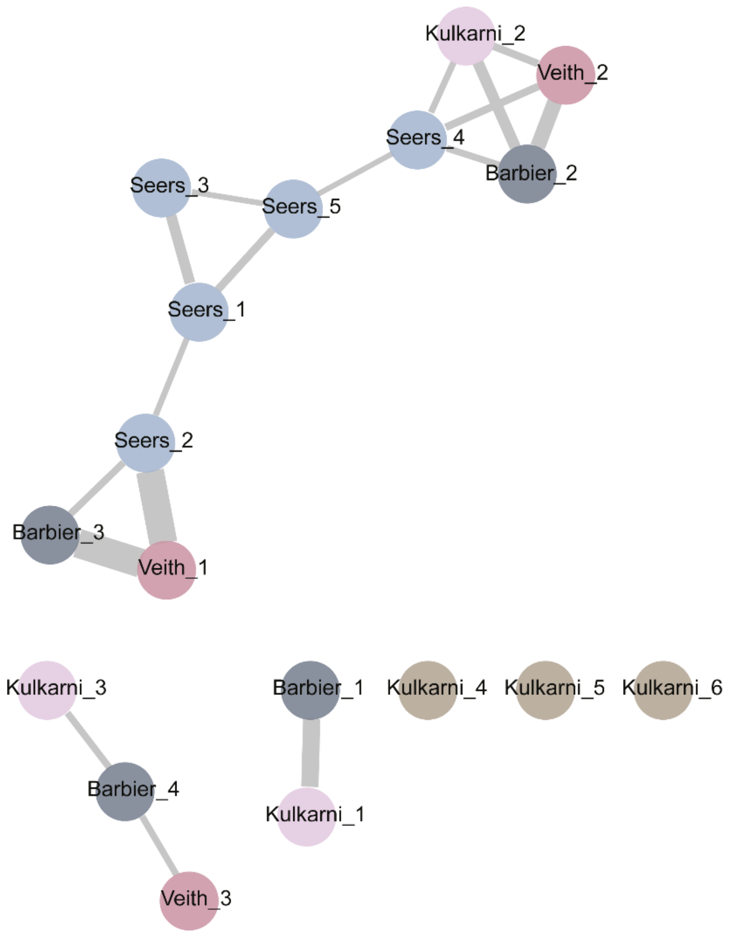


**Figure S3:** Motif similarity network (MSN) of the 18 compiled CTD motifs generated using Tomtom software by setting the overlap >= 3 and score >= 2. We visualized the MSN using Cytoscape, where each node represents the motifs and the edges represent the score computed using Tomtom.

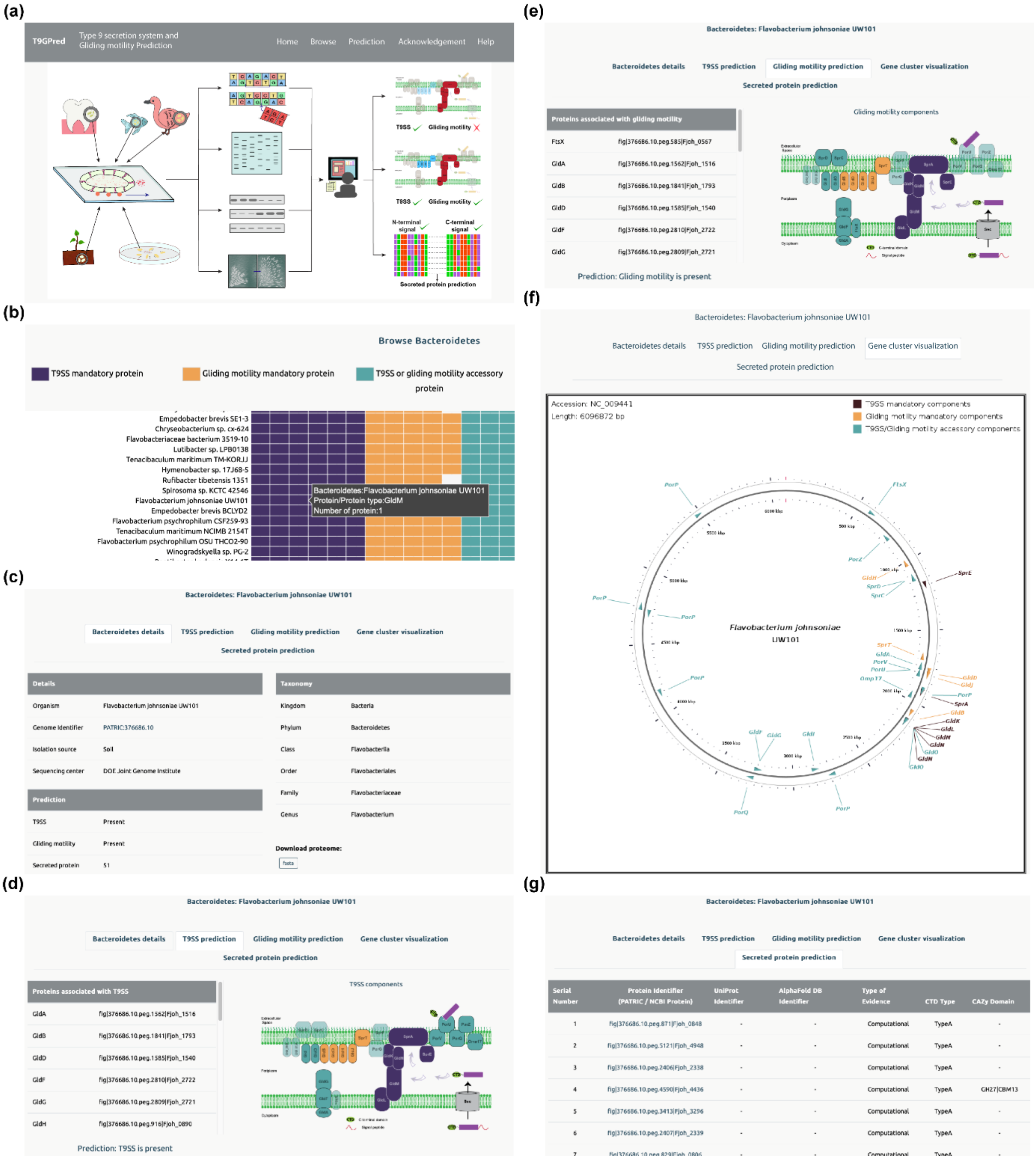

**Figure S4:** Snapshots of the web interface of the T9GPred web server. (a) 'Home' page showing navigation bar. (b) 'Browse' option enables users to explore the details and predictions for 693 completely sequenced *Bacteroidetes* strains considered in this study. (c) The detailed information page for selected *Bacteroidetes* strain. (d) The 'T9SS prediction' page lists the predicted protein components associated with T9SS and visualizes the predicted components. (e) The 'Gliding motility prediction' page lists the predicted protein components associated with gliding motility and visualizes the predicted components. (f) The 'Gene cluster visualization' page provides a visualization of the location of the predicted protein components in the genome of the selected *Bacteroidetes strain*. (g) The 'Secreted protein prediction' page tabulates the proteins predicted to be secreted via T9SS of the selected *Bacteroidetes* strain.

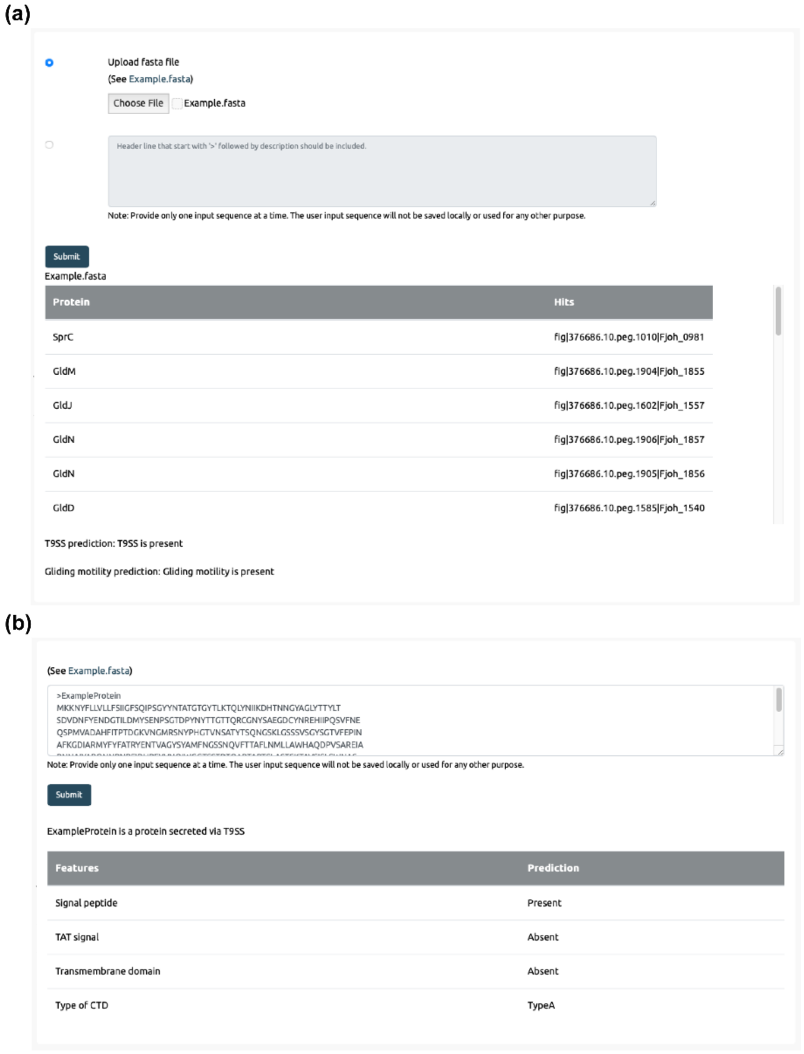


**Figure S5:** Snapshots of the prediction tool in T9GPred web server. (a) The 'Prediction of T9SS and gliding motility' page allows users to submit the proteome of a bacterium and provides the prediction on the T9SS and gliding motility in a tabular format. (b) The 'Prediction of protein secreted via T9SS' page allows users to submit a protein sequence and provides the prediction of the features in a tabular format.

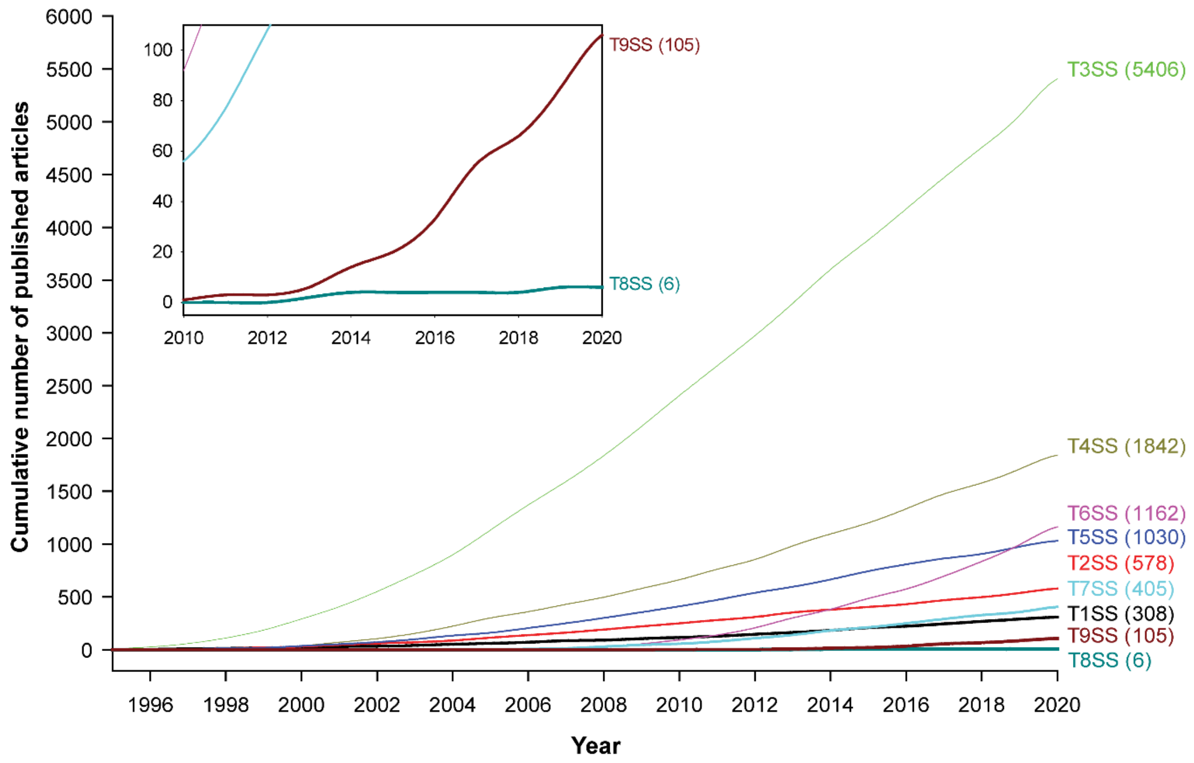


**Figure S6:** Cumulative estimate of published research articles for the nine different secretion systems (T1SS-T9SS) in bacteria. We have not considered T10SS or T11SS in this estimate as these two systems are very recently discovered.

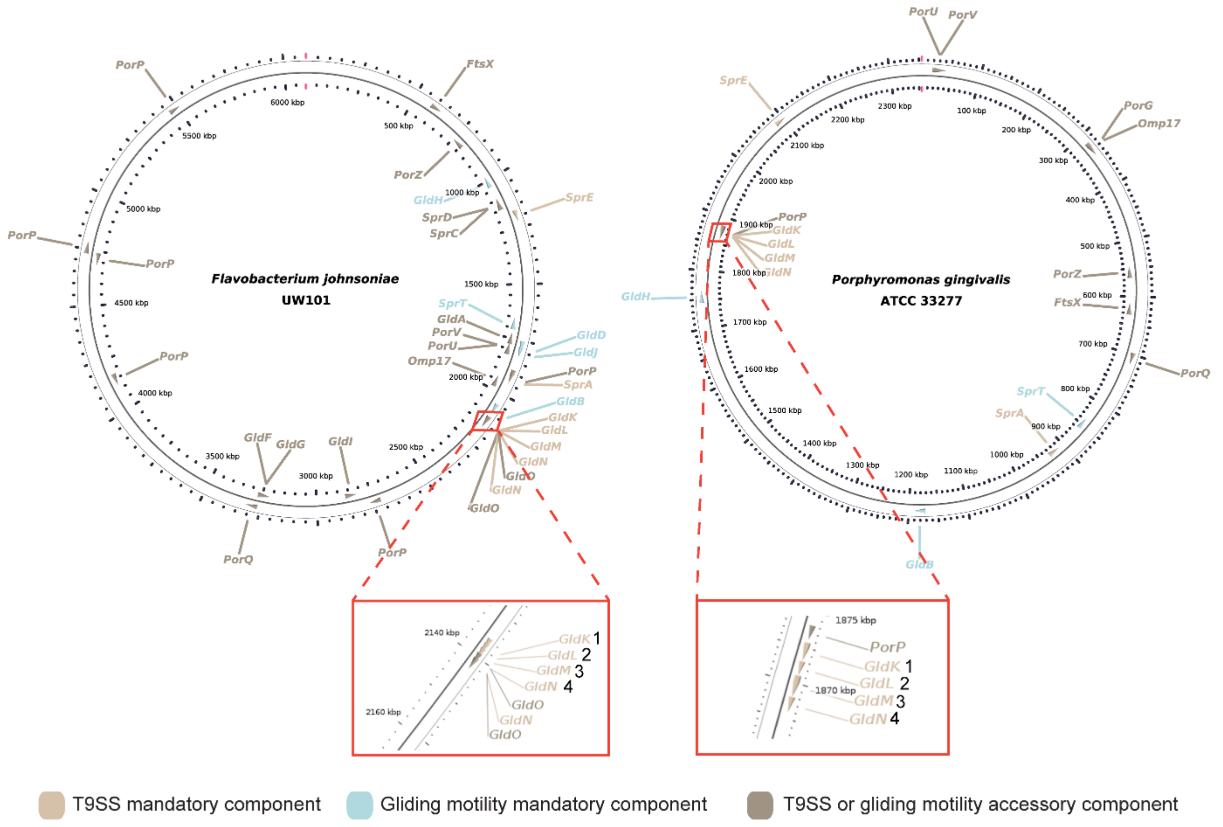


**Figure S7:** Gene cluster visualization of the proteins associated with T9SS or gliding motility in the genome of *Flavobacterium johnsoniae* UW101 and *Porphyromonas gingivalis* ATCC 33277. The operons consisting of GldK (1) GldL (2) GldM (3) GldN (4) are shown in the enlarged view. The figure was generated using CGView package.

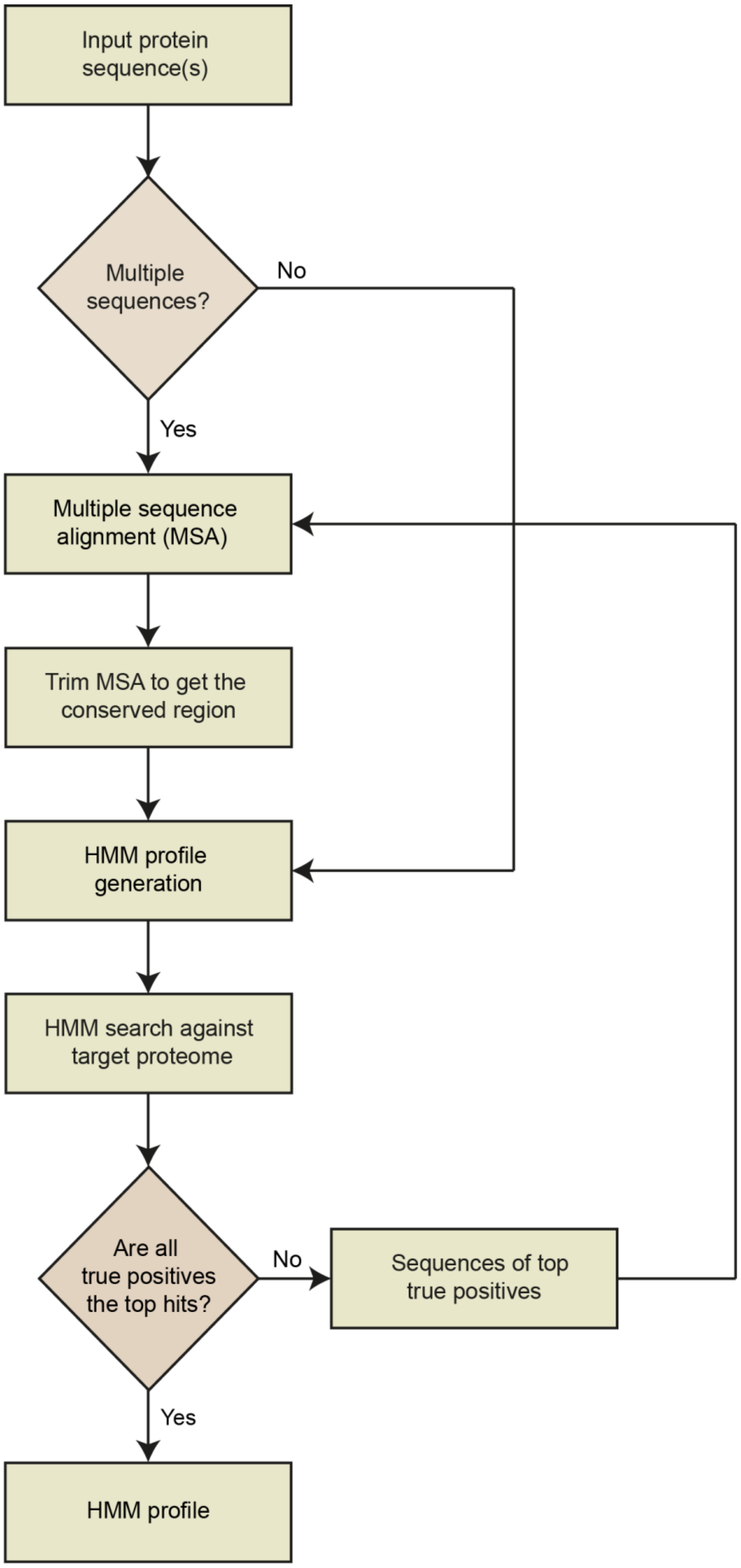


**Figure S8**: Flowchart describing the workflow to create the HMM profile from protein sequences.
